# Chromoplast plastoglobules recruit the carotenoid biosynthetic pathway and contribute to carotenoid accumulation during tomato fruit maturation

**DOI:** 10.1101/2022.09.14.507955

**Authors:** Wayne Zita, Ségolène Bressoud, Gaetan Glauser, Felix Kessler, Venkatasalam Shanmugabalaji

## Abstract

Tomato (*Solanum lycopersicum*) fruit maturation is associated with a developmental transition from chloroplasts (in mature green fruit) to chromoplasts (in red fruit). The hallmark red color of ripe tomatoes is due to carotenogenesis and accumulation of the red carotenoid lycopene inside chromoplasts. Plastoglobules (PG) are lipid droplets in plastids that are involved in diverse lipid metabolic pathways. In tomato, information on the possible role of PG in carotogenesis and the PG proteome is largely lacking. Here, we outline the role of PG in carotenogenesis giving particular attention to tomato fruit PG proteomes and metabolomes. The proteome analysis revealed the presence of PG-typical FBNs, ABC1K-like kinases, and metabolic enzymes, and those were decreased in the PG of tomato chromoplasts compared to chloroplasts. Notably, the complete β-carotene biosynthesis pathway was recruited to chromoplast PG, and the enzymes PHYTOENE SYNTHASE 1 (PSY-1), PHYTOENE DESATURASE (PDS), ZETA-CAROTENE DESATURASE (ZDS), and CAROTENOID ISOMERASE (CRTISO) were enriched up to twelvefold compared to chloroplast PG. We profiled the carotenoid and prenyl lipid changes in PG during the chloroplast to chromoplast transition and demonstrated large increases of lycopene and β-carotene in chromoplast PG. The PG proteome and metabolome are subject to extensive remodeling resulting in high accumulation of lycopene during the chloroplast-to-chromoplast transition. Overall, the results indicate that PGs contribute to carotenoid accumulation during tomato fruit maturation and suggest that they do so by functioning as a biosynthetic platform for carotenogenesis.

## Introduction

Chloroplasts, specialized photosynthetic plastids are present in all green tissues of plants, and are characterized by the presence of an extensive membrane system called thylakoids [1]. Plastoglobules (PG) bud from the external leaflet of the thylakoid membrane and remain associated with the thylakoid membranes [2]. These “osmophilic globuli” were discovered by Lichtenthaler in the early 1970s with the advances in transmission electron microscopy [3]. PG participate in thylakoid lipid metabolism in response to biotic and abiotic stresses (light stress and nitrogen deprivation) [4-6]. Metabolites stored in PG may be exchanged with the thylakoid membrane [7]. PG have dynamic properties changing in number and size and depending on developmental stage and environmental conditions to assure thylakoid homeostasis [8, 9].

Plastoglobules contain a small specialized proteome, consisting of approximately 30 proteins. The components of the Arabidopsis and *Capsicum annuum* (bell pepper) PG proteomes have been categorized into structural proteins, regulatory kinases, and enzymes. Structural proteins named plastid lipid-associated proteins, plastoglobulins or fibrillins (FBNs) have been identified in the proteomes of red bell pepper and Arabidopsis PG [10-12]. The presence of a lipocalin domain suggests that the FBNs may contribute to the transport of lipids or the channeling of metabolites contained in PG [13, 14]. The PG proteomes contain presumed regulatory proteins namely ABC1K-like kinases. ABC1K1 together with ABC1K3 may contribute to plastoquinone distribution within the chloroplast [15, 16]. Together with the FBNs, ABC1Ks are the most abundant components of the PG proteomes [12]. In addition, PGs contain several uncharacterized predicted enzymes but also some well-known ones such as TOCOPHEROL CYCLASE (VTE1) and NAD(P)H DEHYDROGENASE C1 (NDC1) [10, 17]. They are involved in α-tocopherol and phylloquinone biosynthesis, respectively, and implicate PG in these biosynthetic pathways. A considerable number of studies identify PG as a storage compartment containing neutral and prenyl lipids, including phylloquinone (vitamin k), plastoquinone (PQ9), plastochromanol 8 (PC-8), tocopherols, triacylglycerols, fatty acids, and carotenoids [18-21].

In tomato fruit, chromoplasts develop from chloroplasts, a process that involves the dismantling of thylakoid membranes and chlorophyll breakdown as well as the recycling of the breakdown products [22-24]. PG in chromoplasts contain carotenoids such as β-carotene and lycopene, a high level of which is responsible for the hallmark red color of tomato fruit [25]. Tomato chromoplast PG are globular, whereas those of red bell pepper are fibrillar [21, 26, 27]. Fibrillins were first discovered as a major protein component of fibrillar PG in bell pepper, hence their name [21]. Later, red bell pepper chromoplast PG were shown to contain several carotenoid biosynthetic pathway enzymes including LYCOPENE β-CYCLASE (LCYB) and ζ-CAROTENE DESATURASE (ZDS) [11]. To characterize and establish tomato plastoglobule proteomes, we isolated PG from lysed chloroplasts and chromoplasts by sucrose density gradient centrifugation followed by nano LC-MS/MS analysis (Nanoscale liquid chromatography coupled to tandem mass spectrometry) of the associated proteins. We identified multiple known as well as new candidate PG proteins. Notably, enzymes of the carotenoid biosynthetic pathway were recruited to PG and some of them were strongly enriched in chromoplast PG. This coincided with strongly increased accumulation of lycopene supporting a possible role of PG in carotogenesis during tomato fruit ripening and chloroplast to chromoplast differentiation.

## Material & methods

### Plant material and growth conditions

Tomato used was (Solanum Lycopersicum, cv. Micro-Tom). The plants were grown in soil under 200 µmol.m^-2^.s^-1^ in a growth chamber, with a photoperiod of 16 h of day and 8 h of the dark at at 22° and 18°C respectively. The relative air humidity of the growth chamber was around 30%. Fruits were harvested 10 days after the breaker stage and immediately put on ice for further experimentation.

### Fractionation and isolation of tomato chloroplast and chromoplasts plastoglobules

Fruits were washed with distilled water. The peduncle, the gel, and the seeds were removed from the fruits, then the pericarp was cut into small pieces and stored at 4 °C overnight. 130 g of tomato pieces were put into a cold waring blender, 150 ml extraction buffer (0.4 M sucrose, 50 mM tris, 1 mM EDTA, 1 mM DTT) was add before the experiment, pH= 7.8) was added followed by homogenization (twice for 3 seconds at low speed, then twice for 5 seconds at high speed). The homogenate was filtered through three to four layers of cheesecloth (gauze) and put in a 500 ml centrifuge tube on ice. Tubes were centrifuged at 5000x g for 10 min in a Sorvall RC-5B (SLA-1500 Super Lite rotor) at 4 °C. The supernatant was removed, and the pellet was resuspended in 5 ml of extraction buffer and transferred to a 50 ml centrifuge tube on ice. Tubes were centrifuged at 9000x g for 10 min in a Sorvall RC-5B (SM-24 rotor) at 4 °C. The supernatant was removed, and the pellet kept on ice. The pellet was resuspended in 5 ml of 45% sucrose (45 % sucrose [w/v], 50 mM tricine, 5 mM sodium bisulphite, 2 mM EDTA, 2mM DTT add before use, pH7.9). Chromoplasts were homogenized mechanically using a handheld Potter homogenizer. 3 ml of 45 % sucrose were added to the potter homogenizer and 8 ml of the resulting homogenate were transferred to a 38.5 ml Ultra-Clear centrifuge tube. Layers of 38 %, 20 %, 15 % and 5 % sucrose were added, and the gradients centrifuged 20 h at 4 °C at 100 000x g. Western blot analysis indicated that PG were contained in the top sucrose layers.

### Proteins precipitation and immunoblotting

Proteins contained in PG and other fractions were precipitated with acetone. Proteins were separated by SDS-PAGE and transferred to nitrocellulose membrane. The immunoblotting was performed using FBN1A [10], TOC75 [28], and LHCB2 (Agrisera) antibodies.

### Protein identification by nano-LC-MS/MS

The dry pellet of the PG protein sample was resuspended in 45 µl of miST buffer (1% sodium deoxycholate, 100mM Tris pH 8.6 (−20°C), 10 mM DTT, 0.2µM EDTA), vortexed and heated at 90°C for 5 minutes. 160 mM chloroacetamide in 10 mM Tris pH 8.6 was added to the sample and incubated for 30 minutes at room temperature. Next, the tryptic digestion was performed with 0.3ug LysC/Trypsin for 90 minutes at 37C. An OASIS plate was pre-equilibrated with acetonitrile (MeCN) and SCX buffer. 300 µl of 100% ethyl-acetate and 1% trifluoroacetic acid (TFA) were added to the samples, vortexed for 2 minutes, and centrifuged at 5000 rpm for 5 minutes. The bottom aqueous phase was transferred into prefilled SOLA SCX columns and spun through the column (not too fast, typically, 2000 rpm 1 min is sufficient). The column was washed with 300ul ethyl acetate, 0.5% TFA and HPLC solvent A (2% MeCN, 0.1% formic acid). The sample was eluted with 200ul elution buffer (80% MeCN, 19% water, 1% NH3). The eluted sample was dried in a speed vac and resuspended in HPLC solvent A for MS analysis (Fusion, IT mode). Scaffold viewer Scaffold (version Scaffold_5.1.2, Proteome Software Inc., Portland, OR) software was used to validate MS/MS based peptide and protein identifications. Peptide identifications were accepted if they could be established at greater than 90.0 % probability by the Scaffold Local FDR algorithm. All MS/MS spectras were analyzed using Mascot (Matrix Science, London, UK; version 2.6.2) to find the known protein sequences with a fragment ion mass tolerance of 0.50 Da and a parent ion tolerance of 10.0 PPM.

### Prenyl lipid and carotenoid analysis from whole tomato fruit and PG fractions

Prenyl lipids and carotenoids were extracted from whole tomato fruit and PG fractions using established methods [29, 30]. The prenyl quinones and carotenoids were separated and quantified by reverse-phase ultra-high-pressure liquid chromatography coupled to quadrupole-time-of-flight mass spectrometry (UHPLC-QTOFMS). Absolute concentrations of prenylquinones (PQ-9, PC-8, phylloquinone, α-T, γ-T, δ-T, PC-OH, PQH2-9 PQ-OH) and carotenoids (lycopene, lutein, β-carotene, phytoene) were calculated based on standard calibration curves. In addition, the carotenoids violaxanthin and neoxanthin were measured based on lutein standards.

## Results

### Isolation and quality assessment of chloroplast and chromoplast plastoglobules

To determine tomato fruit PG proteomes, tomato plants (*Solanum Lycopersicum*) were grown under standard light conditions (200 µmol.m^-2^.s^-1^) and mature green and ripe red fruit (breaker stage: BS +10) were harvested [31]. PG from chloroplasts (in mature green fruit) and chromoplasts (in red fruit) were isolated by flotation centrifugation on a discontinuous sucrose gradient [26]. The low density of PG allowed separation from plastid membrane compartments by differential flotation. Visual inspection suggested that chloroplast plastoglobules (PG), EN (envelope) and TM (thylakoid membranes) from chloroplasts had been separated well (Fig 1A). The same held true for chromoplast plastoglobules (PG) that also appeared well separated from carotenoid membranous crystals (CR) and envelopes (EN) (Fig 1C). This was assessed by western blotting analysis of the gradient fractions using specific antibodies against marker proteins for PG FBN1A, envelopes TOC75, and thylakoid membranes LHCB2 (Fig. 1 B, D). The established PG marker FBN1A was detected in the first ten low-density fractions but also the last fractions, which has already been shown in previous studies and is likely due to the association of PG with plastid membranes [10, 12, 26](Fig 1B, D). The absence of the thylakoid LHCB2 and the outer membrane TOC75 markers in the low-density fractions indicated PG enrichment. Fractions one to five enriched in PG were selected for further analysis, pooled, and proteins precipitated by acetone. The protein pellets were dried. Two independent biological replicates for both chloroplast and chromoplast PG were obtained.

**Fig 1.**
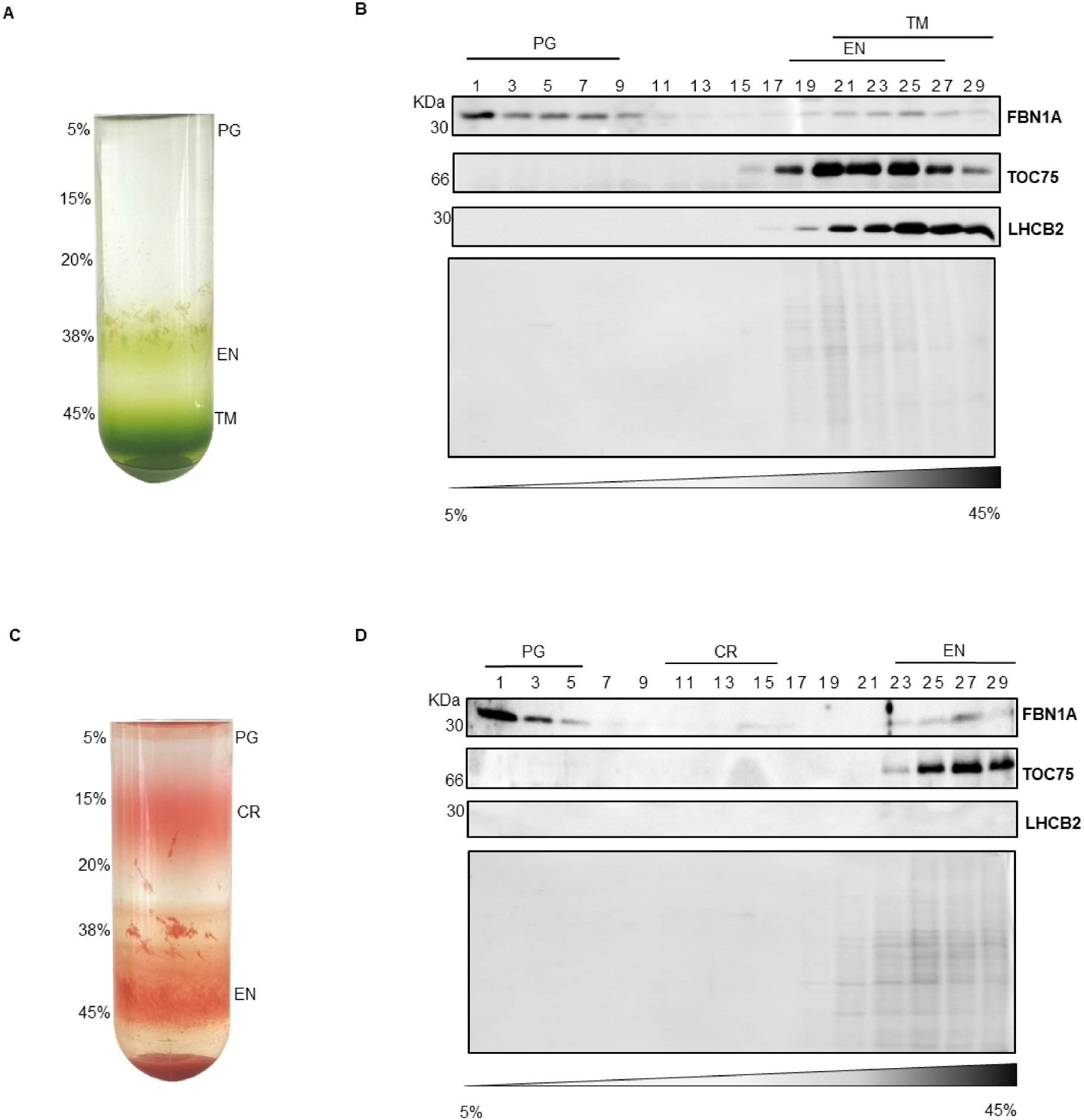
Isolation of tomato fruit PG from chloroplast and chromoplast. (A) Isolated chloroplasts were fractionated on a sucrose step gradient (5, 15, 20, 38, 45% sucrose). PG (plastoglobules); EN (envelope); TM (thylakoid membranes). (B) Uneven fractions from 1 to 29 were subjected to SDS-PAGE followed by immunoblotting using antibodies against FBN1A (PG marker), TOC75 (envelope marker), and LHCB2 (thylakoid marker). (C) Isolated chromoplasts from red tomato fruit were fractionated on a sucrose step gradient (5, 15, 20, 38, 45% sucrose). PG (plastoglobules); CR (carotenoid crystals); EN (envelope). (D) Uneven fractions 1 to 29 were subjected to SDS-PAGE followed by immunoblotting using antibodies FBN1A (PG marker), TOC75 (envelope marker), and LHCB2 (thylakoid marker).

### Tomato plastoglobule proteome

Pooled and precipitated PG fractions were digested with trypsin. The tryptic peptides were analyzed by gel-enhanced liquid chromatography-mass spectrometry (GeLCMS). The data were analyzed using the MASCOT/MaxQuant and Scaffold algorithm to identify proteins corresponding to the peptides. The parameters were set to identify only those individual proteins that were common to both biological replicates of the experiment. A total of 1150 and 838 proteins were identified in chloroplast and chromoplast PG, respectively. These sets of proteins were further analyzed using the UniProt, TAIR and SUBA online resources as well as the TargetP algorithm. This analysis reduced the number to 271 and 147 candidate proteins for chloroplast and chromoplast PG. The established PG proteomes from *Arabidopsis thaliana* and red bell pepper [10-12] combined together were used as references to identify core components of the chloroplast and chromoplast PG proteomes. Thereby, 33 tomato chloroplast and 31 chromoplast proteins were found in common with Arabidopsis, and bell pepper and constituted the tomato fruit PG core proteomes. With the exception of the absence of CCD4 and NDC1 from chromoplast PG, the chloroplast and chromoplast PG core proteomes were identical. Abundant PG proteins including FBNs and predicted regulatory protein kinases ABC1Ks were identified [12]. Also present in both proteomes were prenyl lipid pathway enzymes (including VTE1, PHYTYL ESTER SYNTHASE (PES), ZDS and LCYB) and additional accepted PG proteins (including PGM48, PG18, and SOUL4) [10, 11, 24, 32-34]. In addition, to the known PG proteins, we have identified 18 new candidates for the tomato chloroplast PG proteome and 17 for the chromoplast PG proteome of which 15 were common to both. They were identifed using PPDB and STRING databases as well as localization studies in the literature essentially by exclusion of curated stromal, thylakoid and envelope proteins. This allowed us to identify three additional candidates for carotenoid biosynthesis pathway (PSY1, PDS, CRTISO), for phylloquinone biosynthesis (MenG), for lipid metabolism (LOXC) and for isopentenyl diphosphate biosynthesis (1-deoxy-D-xylulose-5-phosphate synthase 1; DXS1) (Fig 2 and Table 1).

**Fig 2.**
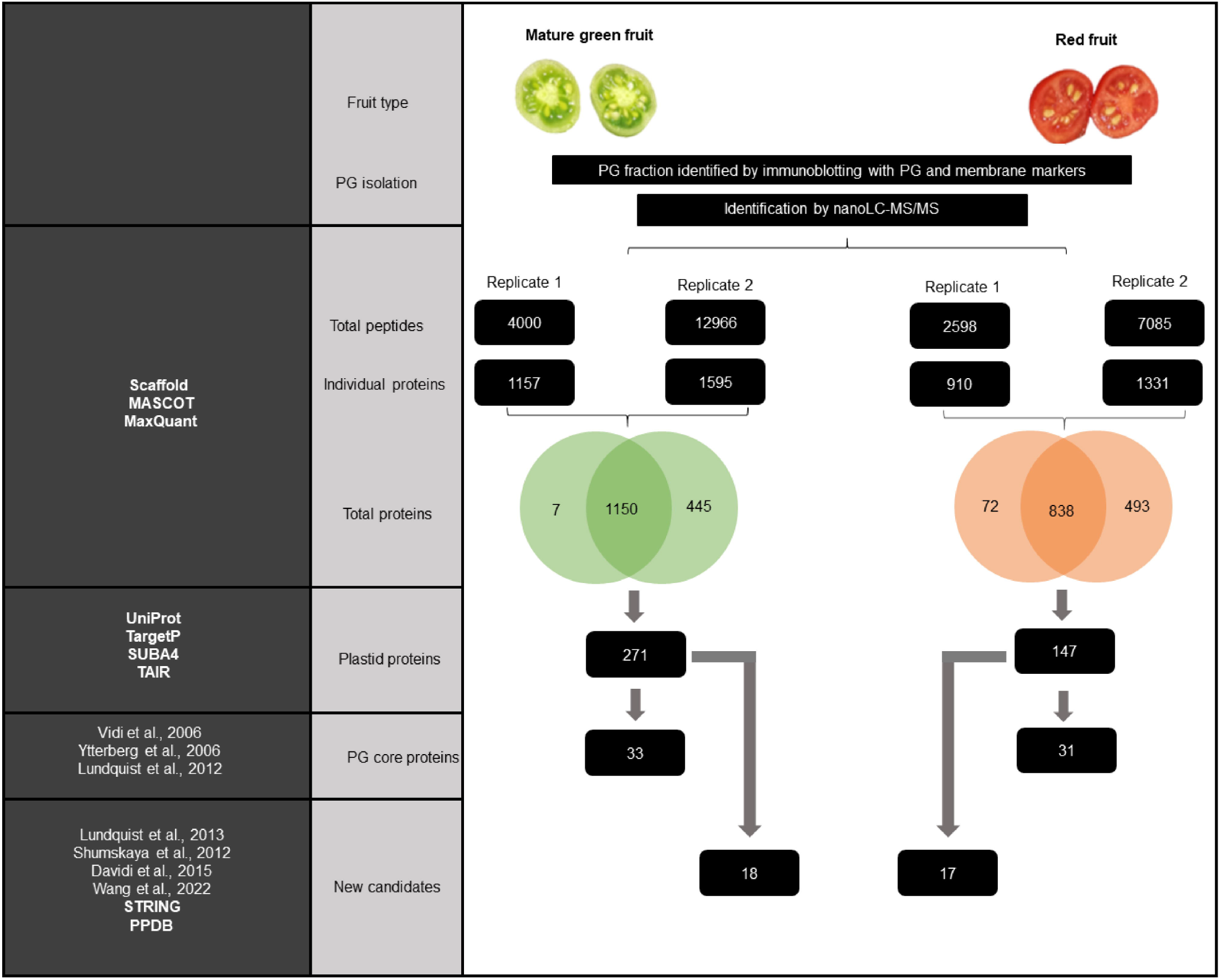
Experimental design used for tomato fruit PG proteome analysis. PGs from mature green and red fruit were isolated separately by sucrose gradient flotation. The two independent replicate PG fractions from mature green and red fruit were analyzed by nano-liquid chromatography (nanoLC)-electrospray ionization (ESI)-tandem mass spectrometry (MS / MS) for peptide identification. All identified peptides (Total Peptides) were further processed by Scaffold, MASCOT, and MaxQuant software to obtain corresponding individual proteins (Individual proteins). The number of individual proteins collected after merging two independent replicates (Total proteins). The Total proteins were filtered using the Uniport, TargetP, SUBA4, and TAIR databases to identify plastid protein (Plastid protein). Plastid proteins were filtered using known chloroplast and chromoplast proteome based on the existing literatures resulting in PG core proteins (PG core proteins). The new PG protein candidates (New candidates) were identified by exclusion of curated stromal, thylakoid and envelope proteins from plastid proteins (Plastid proteins) using PPDB and STRING databases as well as localization studies in the literature.

**Table 1.**
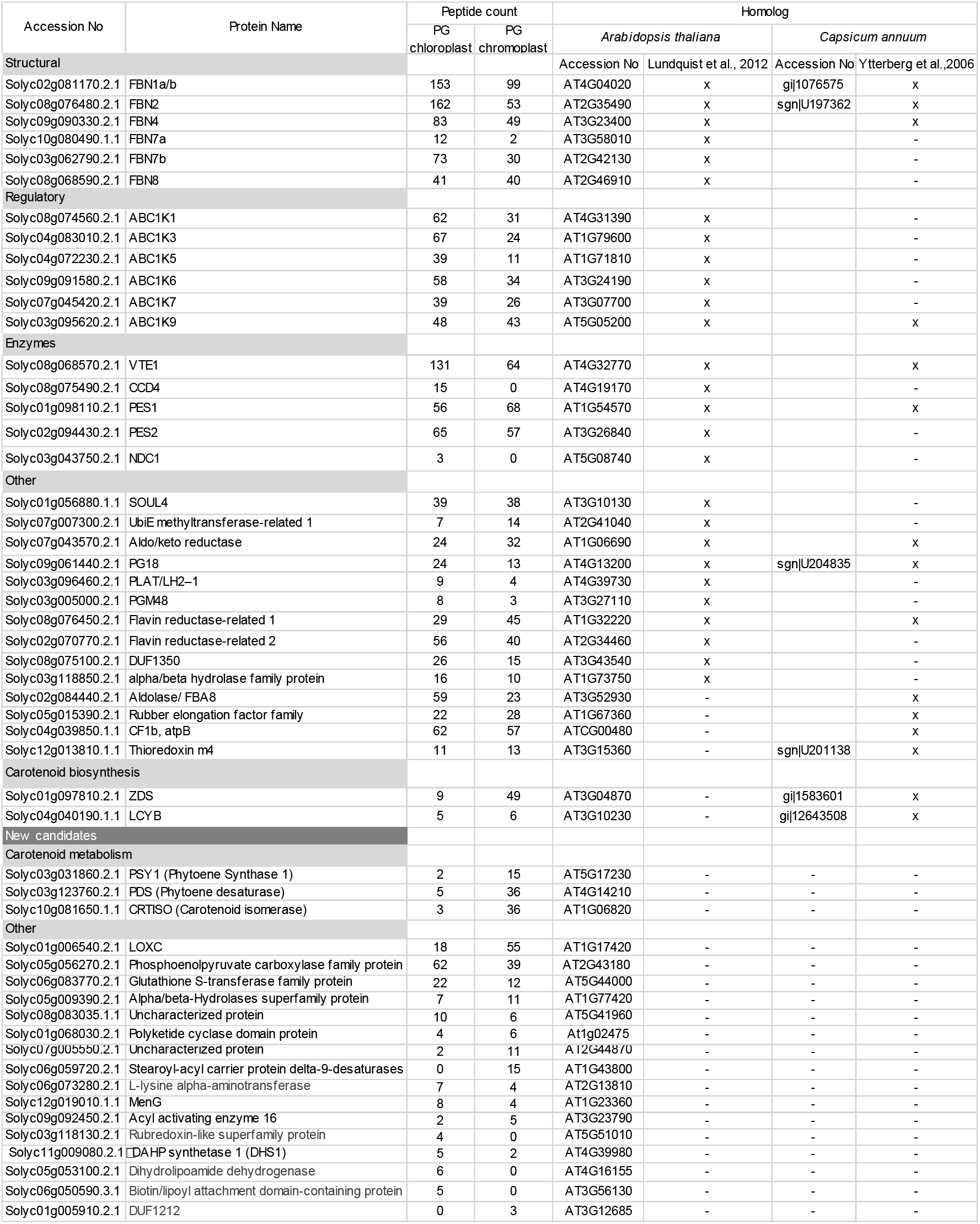
Proteome of tomato fruit chloroplast and chromoplast plastoglobules. Tomato PG proteome which is based on two biological replicates. One representative proteome was selected for the table. The Accession No is the ID identified for each protein on www.solgenomics.net; the common protein name is based on the literature. The accession numbers of homologs are also given based on TAIR for Arabidopsis and Solanaceae database (www.sgn.cornell.edu) for bell pepper. Peptide count represents the number of peptides detected in each sample. The known “core” PG proteome was categorized into five groups and the new candidate PG protein group into two categories.

### Carotenoid biosynthetic enzymes are enriched in chromoplast PG

Five enzymes of the carotenoid biosynthesis pathway upto β-carotene were associated with chloroplast as well as chromoplast PG. The comparison of chloroplast with chromoplast PG revealed differences in their peptide counts. The heatmap based on peptide counts shows a clear increase in chromoplast PG of the first four enzymes required for lycopene biosynthesis namely PSY1, PDS, ZDS, and CRTISO. The higher numbers likely reflect higher enzyme levels that underpin lycopene biosynthetic activity in the chromoplast and during the chloroplast-to-chromoplast transition (Fig. 3). Peptide counts for LYCB were similar in both plastid types, suggesting a constant activity of this enzyme in PG during chloroplast to chromoplast conversion (Fig. 3). Heatmaps also showed reduced presence of FBNs and ABC1Ks in the chromoplast PG compared with chloroplast PG (Fig. S1).

**Fig 3.**
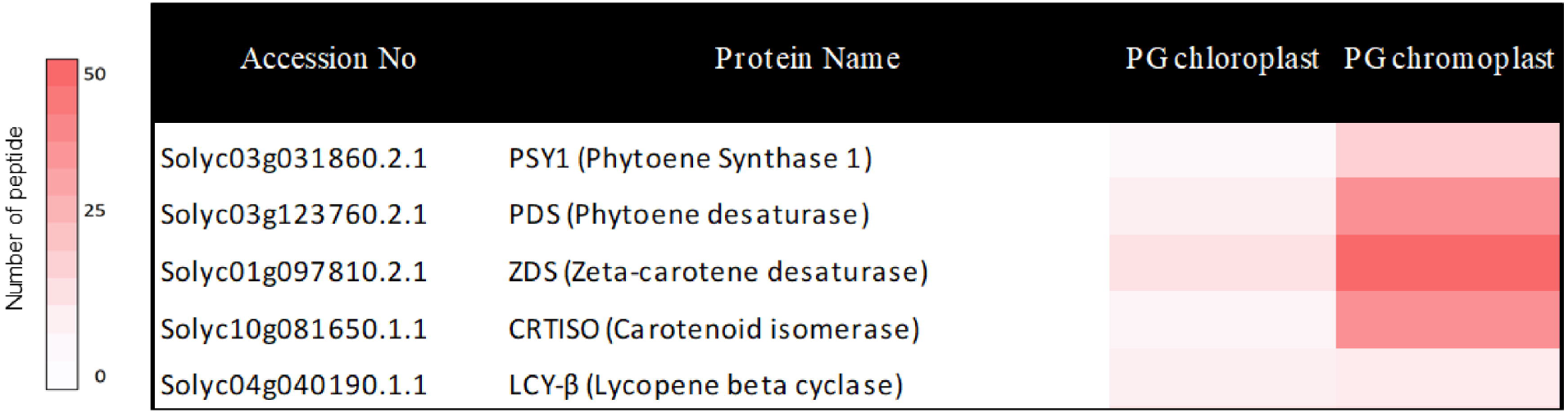
Carotenoid biosynthetic enzymes enriched in tomato PG chromoplasts. The carotenoid biosynthetic enzymes heatmap was generated from peptide counts obtained from PG isolated from chloroplasts and chromoplasts, respectively.

### Protein-protein network of chromoplast PG protein

The chromoplast PG proteome contains various proteins annotated with a large range of functions. We carried out an analysis of the protein network of the chromoplast plastoglobule proteome using STRING (www.string-db.org). The classification and clusters of chromoplast PG proteins proposed by STRING are based on a curated database, experimental results, predicted gene neighborhood, annotated function, text mining, protein co-expression, and protein homology according to the recent literature [35, 36]. Application of STRING resulted in three distinct clusters and a number of non-associated proteins (Fig. 4). Cluster 1 was mainly composed of carotenoid biosynthetic enzymes including PSY1, which condenses two geranylgeranyl diphosphates to phytoene. PDS and ZDS carry out desaturation reactions, which are required for synthesis of 9,9′-di-*cis*-ζ-carotene, and prolycopene, respectively. CRTISO (carotene isomerase) converts prolycopene into all-trans-lycopene. LYCB carries out the cyclization of lycopene into β-carotene [25]. In addition, carotenoid biosynthesis key regulatory enzyme DXS1 [37] was part of cluster 1. Interestingly, FBN1 (CHRC) is part of cluster 1 and also closely associated with cluster 2. FBN1 (CHRC) is well known for its involvement in carotenoid accumulation and fibril formation but also carotenoid stabilization in chromoplasts [21]. Cluster 2 contain FBNs, VTE1, MenG, ABCK1 and ABC1K3. Functions of FBNs have been linked to tolerance to abiotic and biotic stress [4, 5]. Recent studies have shown that ABC1K1 and ABC1K3 are involved in the regulation of photosynthetic adaptation via plastoquinone pool homeostasis. [15, 16]. The tocopherol cyclase (VTE1) catalyzes the formation of the chromanol ring in α-tocopherol biosynthesis and may be regulated via phosphorylation by ABC1K1 [10, 38]. MenG catalyzes the final methylation step of 2-phytyl-1,4-naphthoquinone resulting in phylloquinone [39]. In conclusion, cluster 2 functions are mostly linked to prenyl quinone metabolism. Cluster 3 contains PES, PGM48 and ABC1Ks. PGM48, a PG-associated protease, which in Arabidopsis mainly interacts with other PG proteins and degrades them during senescence [32]. PES belongs to esterase/lipase/thioesterase and acyl transferase family and synthesizes phytol esters and triacyl glycerols during stress and senescence as well as carotenoid esters during the chloroplast-to-chromoplast transition [24]. ABC1K7 regulates the chloroplast membrane remodelling under stress and may act in the oxidative stress resistance pathway [40, 41]. Thus, cluster 3 functions are mostly linked to chloroplast senescence and thylakoid dismantling, processes that also take place during fruit ripening and chloroplast-to-chromoplast transition.

**Fig 4.**
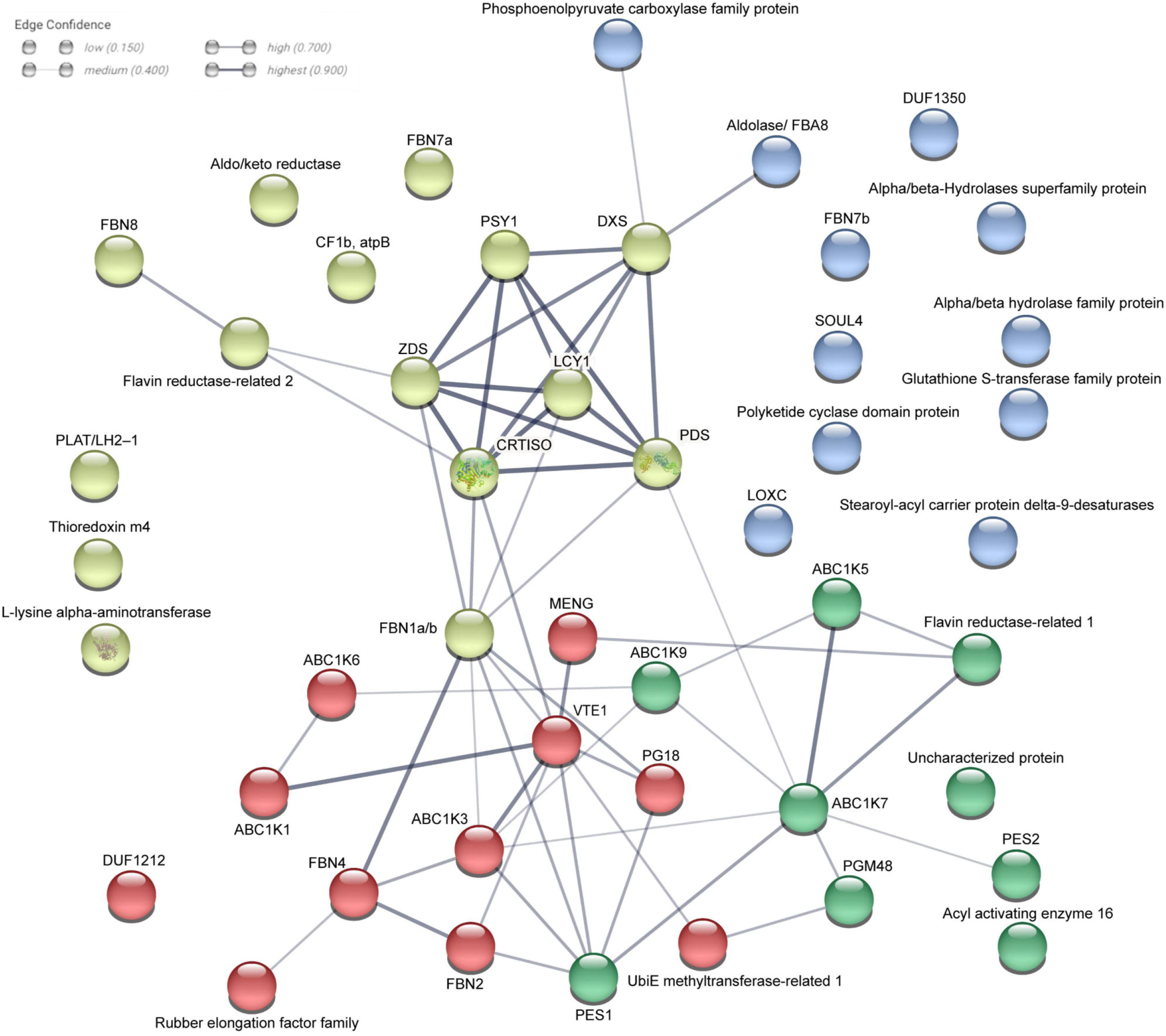
Interaction network of chromoplast PG protei. The tomato chromoplast PG proteome analysis using the STRING software identified three clusters. Pear, (cluster 1) enriched in carotenoid biosynthetic enzymes; Red, (cluster 2) enriched in prenyl quinone metabolism and regulation; green, (cluster 3) enriched in chloroplast senescence and thylakoid membrane dismantling; blue, proteins not associated with the network.

### Lycopene and β-carotene were enriched in the chromoplast PG

We analysed the total carotenoid content and composition in mature green fruit and red fruit as well as in PG purified from chloroplasts and chromoplasts using ultra-HPLC coupled with atmospheric pressure chemical ionization-quadrupole time-of-flight mass spectrometry (UHPLC-APCI-QTOF-MS). Lycopene and β-carotene accumulated to higher concentrations in red fruit than mature green fruit, lycopene being almost undetectable in mature green fruit (Fig. 5A) while lutein and violaxanthin+neoxanthin accumulated to lower concentrations in red fruit than mature green fruit (Fig. S2B). With regard to isolated PG, the relative concentration of lycopene was much higher in chromoplast PG than in chloroplast PG in equal volumes of the isolated fractions (Fig. 5A) while the relative concentrations of lutein were the same and those of β-carotene higher and those of violaxanthin+neoxanthin lower (Fig. 5B). In addition, we compared the relative distribution of lycopene, lutein, β-carotene and phytoene in equal volumes of the isolated fractions of PG, CR (carotenoid crystals) and EN (envelope) fractions. Lycopene and β-carotene levels were highest in CR but in addition to PG the envelopes also accumulated some lycopene and β-carotene (Fig. S3).

**Fig 5.**
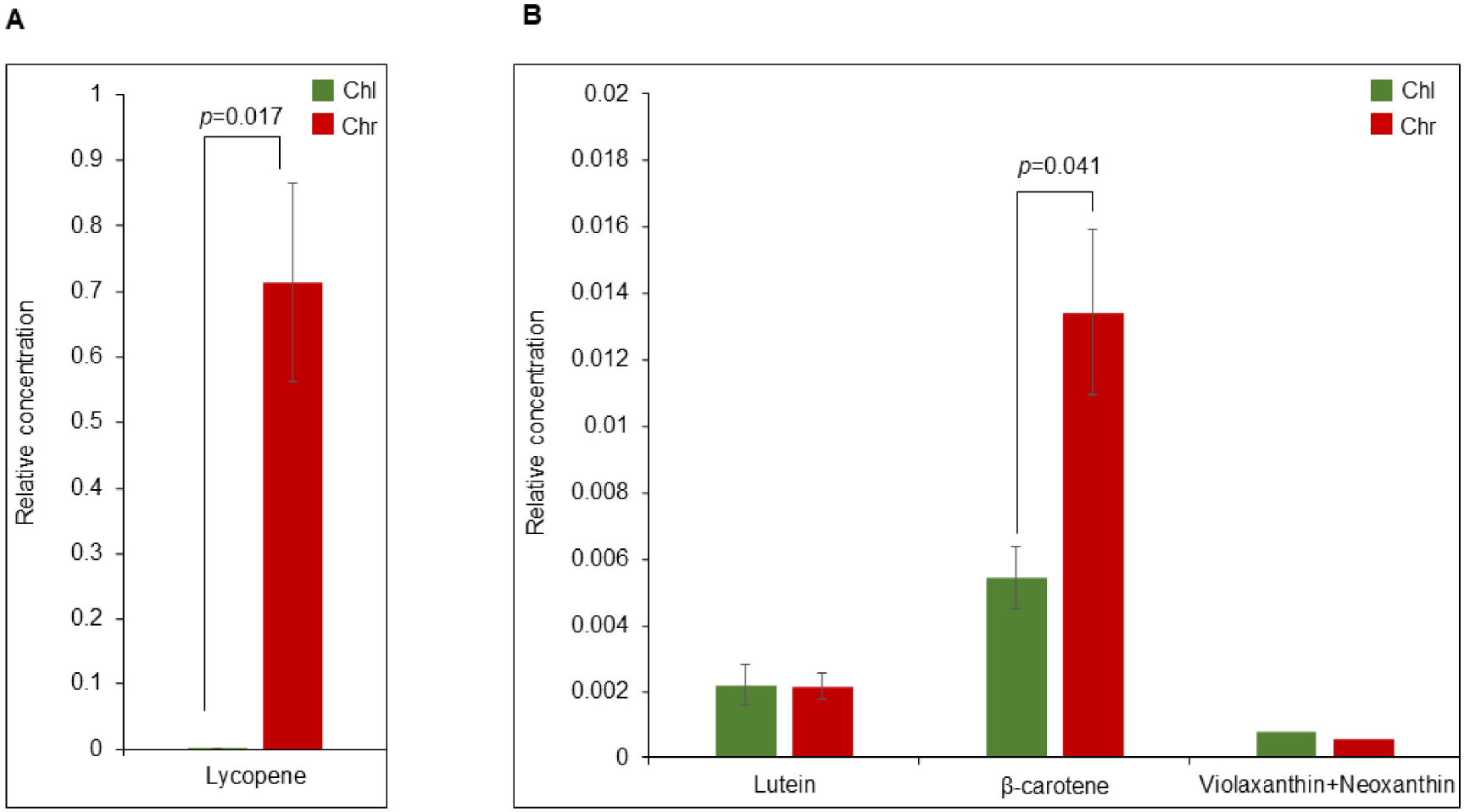
Lycopene and β-Carotene accumulate to high levels in chromoplast PG. (A) Total carotenoids were extracted from equal volumes of gradient fractions containing chloroplast (Chl) or chromoplast (Chr) PG. Lycopene was quantified. (B) Quantification of lutein, β-carotene, and violaxanthin/neoxanthin. All values in the Fig. are the mean of 3 biological replicates (n=3). Statistical differences were assessed with students’ t test and p values are indicated.

### Differential accumulation of prenyl lipids in PG upon chromoplast differentiation

We investigated the total prenyl lipid contents of mature green fruit and red fruit. We obtained prenyl lipid profiles by UHPLC-APCI-QTOF-MS. Red fruit accumulated higher concentrations of plastoquinone (PQ-9), plastochromanol (PC-8), α-tocopherol (α-T), γ-tocopherol (γ-T) and δ-tocopherol (δ-T) while the phylloquinone concentration was lower in red than in mature green fruit (Fig S4). Interestingly this was reversed in PG, the results using equal volumes of the isolated fractions revealing that the relative concentrations of PQ-9 and its derivatives (PC-8, PC-OH, PQH2-9 and PQ-OH), phylloquinone, α-T, γ-T were considerably lower in the chromoplast than in the chloroplast PG (Fig 6).

**Fig 6.**
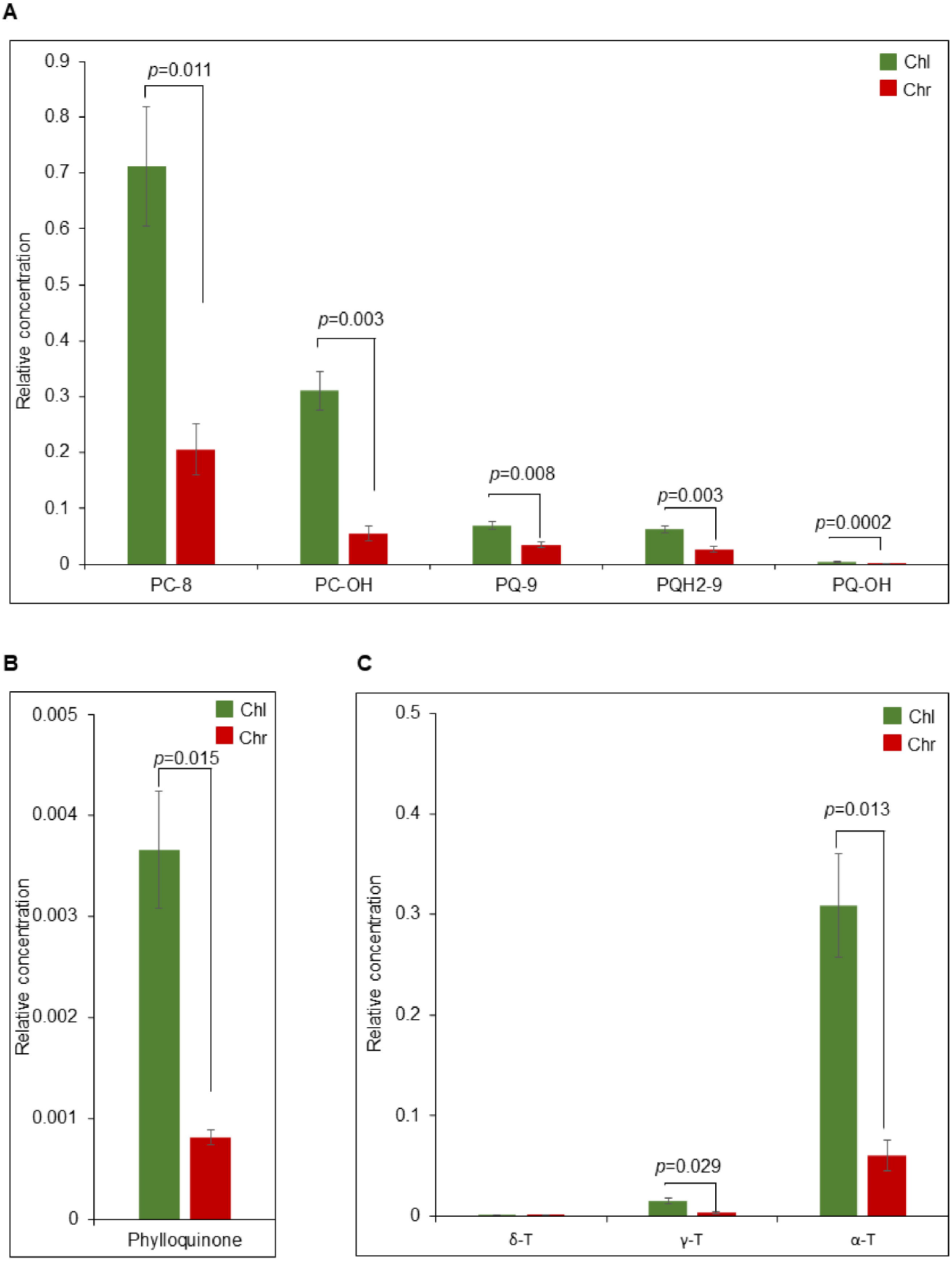
Reduced levels of prenyl quinones in chromoplast PG. (A) The total prenyl quinones were extracted from equal volumes of gradient fractions containing chloroplast (Chl) and chromoplast (Chr) PG, PC-8, plastochromanol; PC-OH, hydroxy-plastochromanol; PQ-9, plastoquinone; PQH_2_-9, plastoquinol and PQ-OH, hydroxy-plastoquinone were quantified (B) Quantification of phylloquinone. (C) Quantification of tocopherols. All values in the Fig. are the mean of 3 biological replicates (n=3). Statistical differences were assessed with student’s t test and p values are indicated.

## Discussion

Purified PG were obtained from the chloroplasts and chromoplasts of mature green and red tomato fruit, respectively, and proteome analysis was carried out resulting in a long list of proteins. UniProt, SUBA4, TAIR database and TargetP algorithm were used to filter the list and identify plastid proteins. Comparison of the resulting list with known PG proteomes indicated that tomato fruit chromoplasts and chloroplast PG together have a total of 33 proteins in common with Arabidopsis and red bell pepper (Table 1). This finding points to the overall functional conservation of PG in different species, tissues and plastid types. These “known” PG proteins are considered core components of the chromoplast PG proteome based on previous reports i.e. CHRC (FBN1A) [21, 42-44]. Western blot analysis confirmed enrichment of CHRC (FBN1A) in PG fractions of chromoplasts and chloroplasts (Fig.1D). In addition to the core components, 17 new candidates that included carotenoid biosynthetic enzymes were identified based on the literature, STRING and PPDB annotation (Table 1) (Fig. 2).

The 17 new candidates of chromoplast PG were attributed to two categories, carotenoid biosynthetic enzymes and “others”. In the “others” category was lipoxygenase C (LOXC), which is specifically expressed during tomato fruit ripening. LOXs participate in fatty acid catabolism during the disassembly of the thylakoid membranes during the chloroplast to chromoplast transition [45]. Moreover, previous studies propose a model in which the C-terminal PLAT (for <u>p</u>olycystin-1, lipoxygenase and <u>a</u>lpha toxin) domain may be responsible for LOX association with PG [46, 47]. Another intriguing candidate is MenG that is involved in phylloquinone biosynthesis [39]. MenG cooperates with the known PG protein NDC1 in the final methylation step of phylloquinone biosynthesis and assignment as PG candidate is therefore not surprising. Moreover, it has been shown previously that fluorescently-tagged MenG and multiprotein of phylloquinone biosynthesis (PHYLLO) resulted in patterns resembling those of PG [48]. Also, in this study, 1-deoxy-D-xylulose-5-phosphate synthase 1 (DXS1), key enzyme of the plastid isoprenoid pathway was present in the tomato PG. A recent report has indicated that both MenG and DXS1 are associated with PG via FBN and PDS in Chlamydomonas [49].

Chromoplast PG contained the carotenoid biosynthesis enzymes ZDS and LYCB as well as three other new candidates PSY1, PDS and CRTISO. Together these enzymes constitute the complete β-carotene biosynthesis pathway downstream of geranylgeranyl diphosphate. Based on the peptide count for PSY1, PDS, ZDS, and CRTISO, they are present at up to 12-fold higher levels in chromoplast PG than in chloroplast PG. However, the LYCB peptide count was almost the same in chromoplast PG and chloroplast PG (Fig. 3; Table 1). The STRING analysis revealed that the enzymes of the carotenoid biosynthesis pathway form a distinct cluster 1 within the chromoplast PG proteome (Fig. 4). Cluster 2 was enriched in enzymes involved in prenyl quinone metabolism and regulation and Cluster 3 in chloroplast senescence- and thylakoid dismantling-associated proteins. Cluster 2 and 3 therefore identify two additional categories of functions that are critical during the chloroplast-to-chromoplast transition. The increased peptide counts of the first four β-carotene biosynthetic enzymes in chromoplast PG compared with chloroplast PG (Fig.3) indicates that PG recruit carotenoid biosynthetic enzymes and may turn them into a biosynthetic platform during carotenogenesis. Our lipidomic study reveals that lycopene and β-carotene became enriched in chromoplast PG during fruit ripening suggesting active contribution to biosynthesis (Fig.5). In agreement with an active role of PG in carotenoid metabolism in tomato, the chromoplast PG enzyme PALE YELLOW PETAL (PYP1) confers yellow flower pigmentation by carotenoid esterification [50]. With regard to PSY1, transient expression experiments provided evidence for PG localization in rice and Arabidopsis [51]. In addition, it was recently reported that bacterial phytoene synthase CrtB localised to PG and increased carotenoid levels when chloroplast-to-chromoplast transition was induced by CrtB expression [52]. Interestingly, a proteome study in the green algae *Dunaliella bardawil* revealed that β-carotene biosynthesis enzymes PDS, ZDS, and CRTISO were present in β-carotene-rich PG [53]. A recent study showed that PDS is one of the main protein components of Chlamydomonas PG [49]. Interestingly, carotenoid cleavage dehydrogenase 4 (CCD4) was identified only in chloroplast PG. PG protease PGM48, a chloroplast PG protease, interacts with CCD4 and degrades it during senescence [32]. Therefore, the presence of PGM48 in chloroplast and chromoplast PG might explain the disappearance of CCD4 during the chloroplast-to-chromoplast transition. We proposed a model for chromoplast PG hosting the carotenoid biosynthetic pathway. In the model, chromoplast PG function as a metabolic platform for carotenoid biosynthesis, especially for lycopene biosynthesis and accumulation (Fig. 7). However, the data also indicate that carotenoid crystals accumulate a large proportion of lycopene and β-carotene (Fig. S3). In summary the data suggest that PG make an important contribution to fruit quality particularly to carotenoid accumulation.

**Fig 7.**
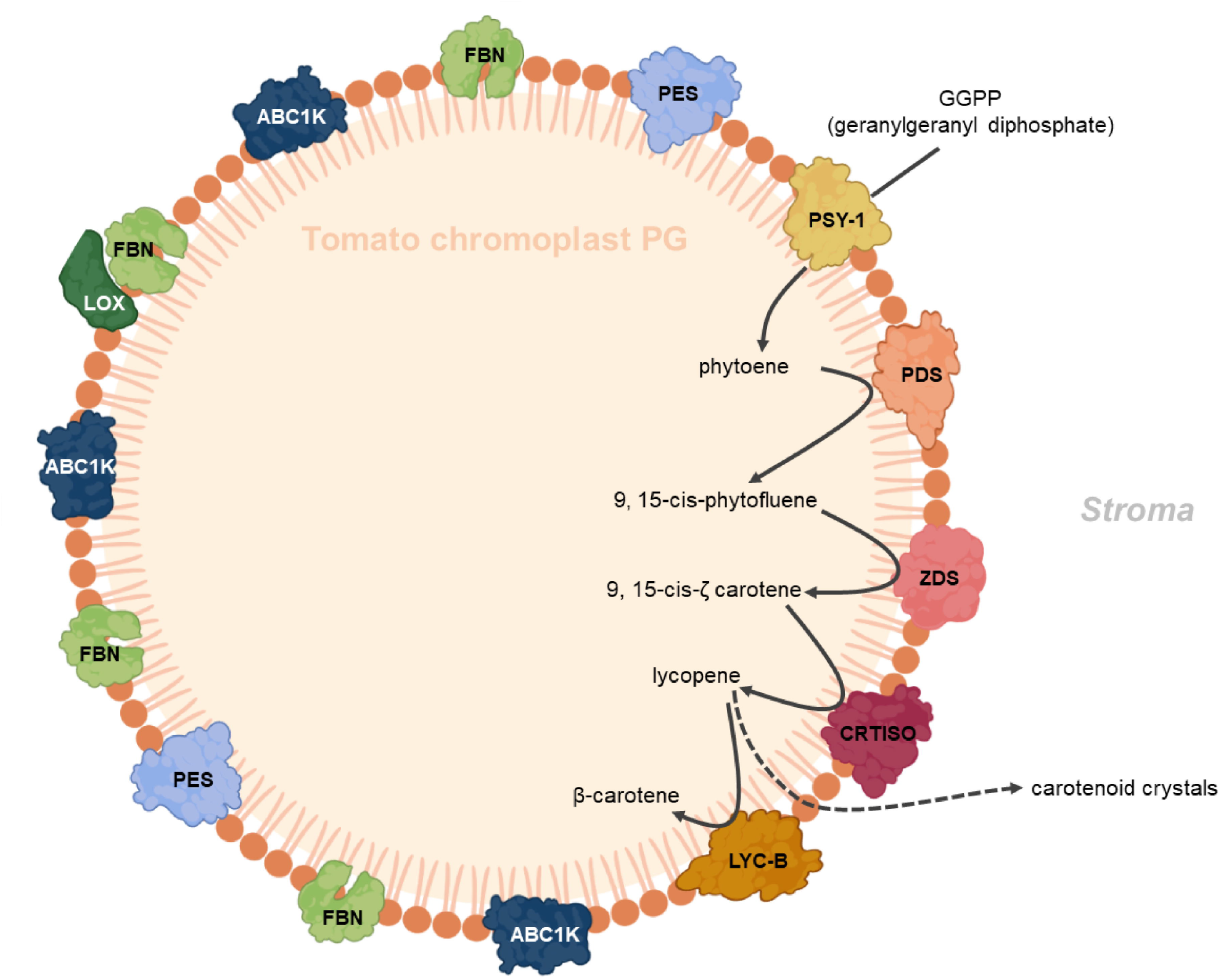
Carotenoid biosynthetic pathway in tomato chromoplast PG. Chromoplast plastoglobules recruit the carotenoid biosynthetic pathway enzymes: phytoene synthase 1 (PSY1), phytoene desaturase (PDS), z-carotene desaturase (ZDS), and carotenoid isomerase CRTISO) and lycopene β-cyclase (LYC-β) to promote carotenoid biosynthesis. Lipoxygenase (LOX) may associate with PG, FBNs have structural functions and may contribute to carotenoid sequestration in PG. Phytyl ester synthase (PES) synthesizes phytyl esters and triacyl glycerol during thylakoid dismantling and Activity of BC1 complex kinase 1 (ABC1K1) is implicated in regulation of prenyl lipid metabolism.

## Author Contributions

WZ, FK, GG and VS designed the experiments. WZ, SB, and GG carried out the experimental work. WZ, FK, GG and VS analysed the data. VS and WZ wrote the manuscript with the help of FK.

## Acknowledgments

We thank the Protein Analysis Facility (PAF), University of Lausanne for supporting mass spectrometry acquisition and analysis.

