## Supplemental Fig 1-4 for "Chromoplast plastoglobules recruit the carotenoid biosynthetic pathway and contribute to carotenoid accumulation during tomato fruit maturation"

### *Supplementary Material*

#### Supplementary Figures

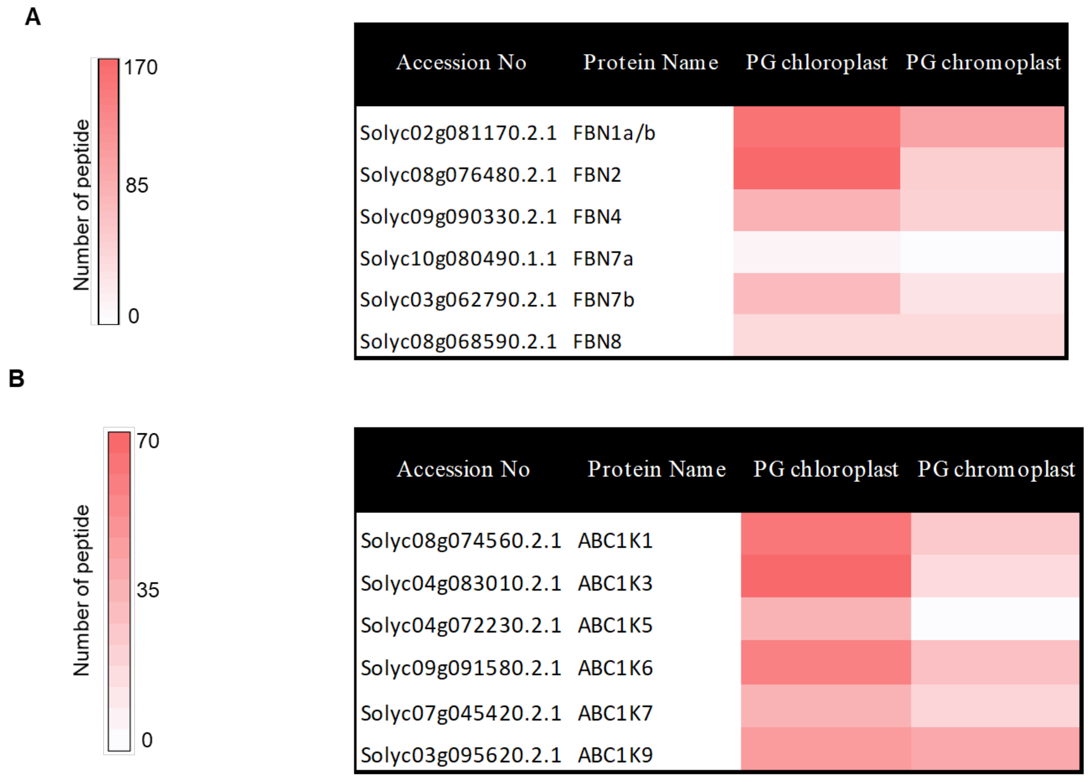


**Figure S1: FBNs and ABC1K- kinases were reduced in tomato chromoplast PG.**

(A) FBN heatmap and (B) ABC1K-like kinase heatmap were generated from peptide counts obtained from PG isolated from chloroplast and chromoplast, respectively.

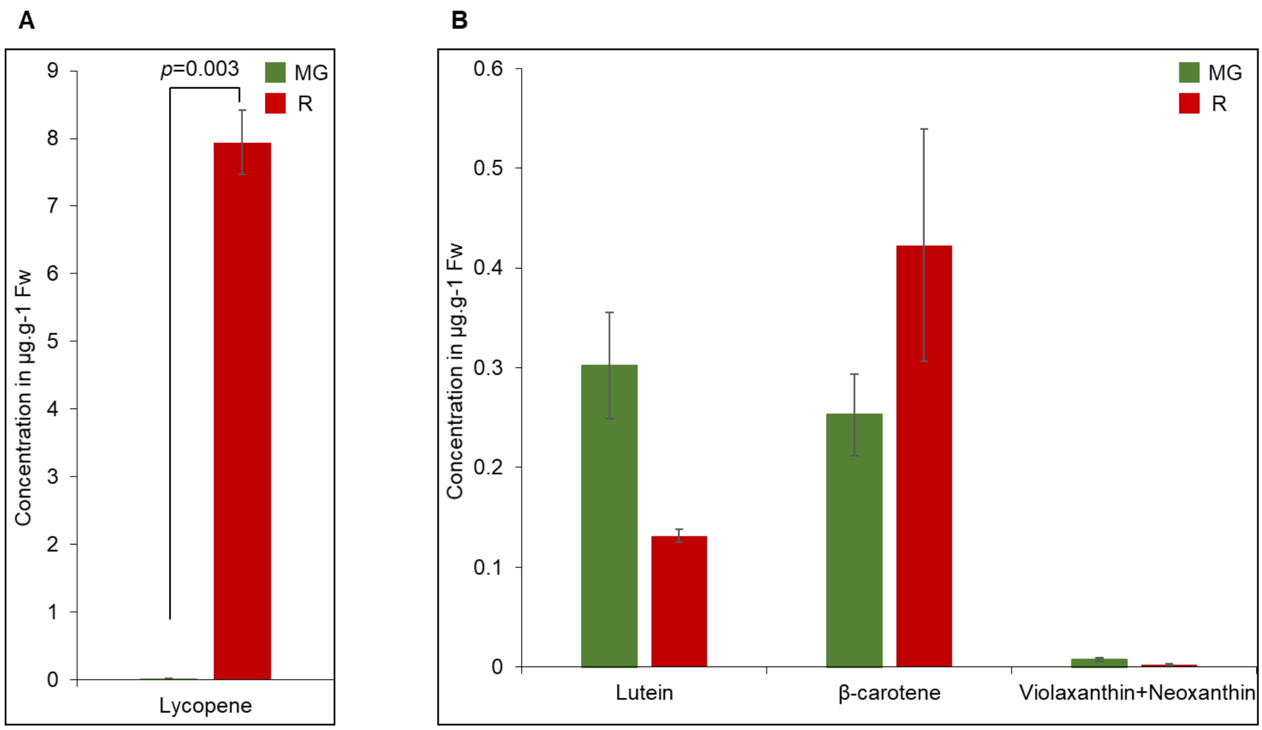


**Figure S2: Lycopene was highly accumulated in the red tomato fruit**

(A) Total carotenoids were extracted from mature green (MG) and red (R) tomato fruit and lycopene was quantified. (B) Quantification of lutein, β-carotene, and violaxanthin/neoxanthin. All values in the figure are the mean of 3 biological replicates (n=3). Statistical differences were assessed with student's t test and p values are indicated.

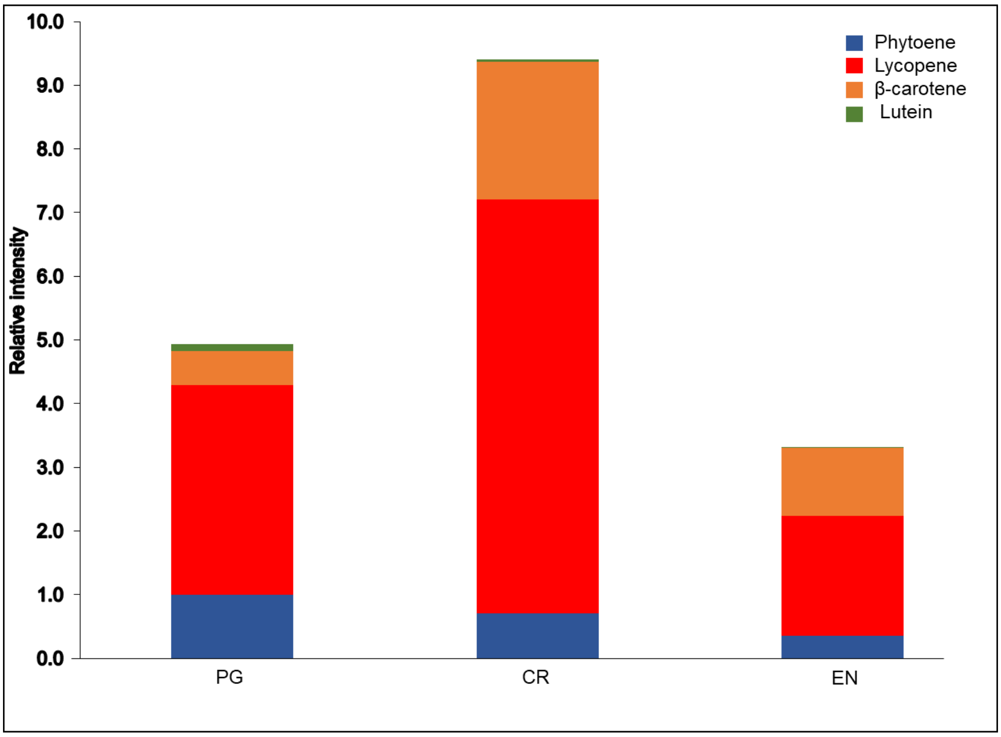


**Figure S3: Carotenoids were differentially accumulated in the chromoplast sub-compartments**

The total carotenoids were extracted from equal volumes of mature red (R) tomato fruit PG (plastoglobules); CR (carotenoid crystals) fractions; EN (envelope). The isolated lycopene, phytoene, β-carotene, and lutein were quantified. Values are the mean of 3 biological replicates (n=3

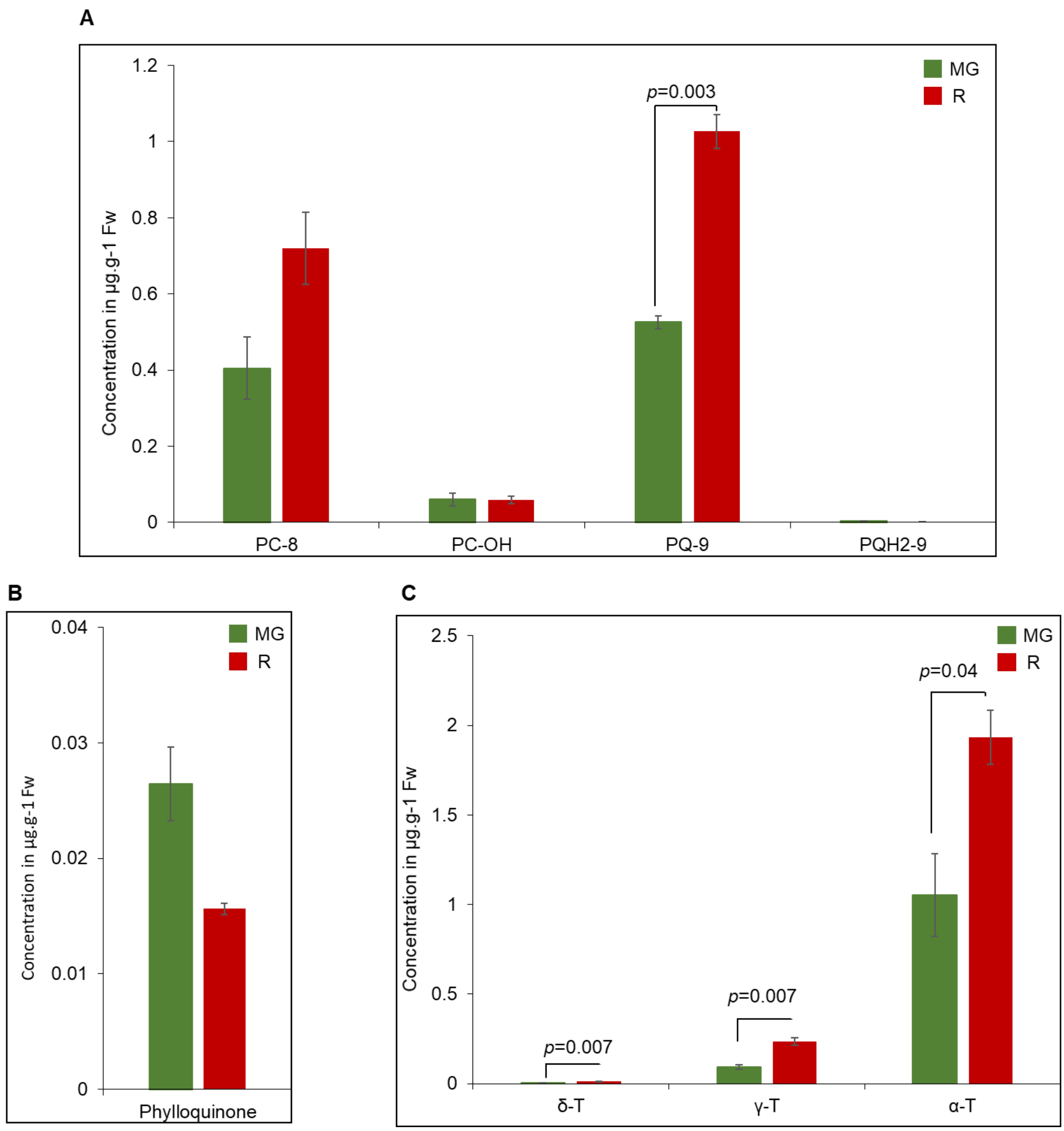


**Figure S4: Tocopherols and plastoquinone were highly accumulated in the red tomato fruit**

(A) The total prenyl quinones were extracted from mature green (MG) and red (R) tomato fruit, PC-8, plastochromanol; PC-OH, hydroxy-plastochromanol; PQ-9, plastoquinone; and $PQH_2$-9, plastoquinol were quantified (B) Quantification of phylloquinone. (C) Quantification of tocopherols. All values in the figure are the mean of 3 biological replicates (n=3). Statistical differences were assessed with student's t test and p values are indicated.
